# Uptake without inactivation of human adenovirus type 2 by *Tetrahymena pyriformis* ciliates

**DOI:** 10.1101/2023.04.13.536677

**Authors:** Margot Olive, Jean Daraspe, Christel Genoud, Tamar Kohn

**Affiliations:** Laboratory of Environmental Chemistry, School of Architecture, Civil and Environmental Engineering (ENAC), Ecole Polytechnique Fédérale de Lausanne (EPFL), Lausanne, Switzerland; Electron Microscopy Facility, Faculty of Biology and Medicine, University of Lausanne (UNIL), Lausanne, Switzerland

## Abstract

Human adenoviruses are ubiquitous contaminants of surface water. Indigenous protists may interact with adenoviruses and contribute to their removal from the water column, though the associated kinetics and mechanisms differ between protist species. In this work, we investigated the interaction of human adenovirus type 2 (HAdV2) with the ciliate *Tetrahymena pyriformis*. In co-incubation experiments in a freshwater matrix, *T. pyriformis* was found to efficiently remove HAdV2, with ≥ 4 log_10_ removal over 72 hours. Neither sorption onto the ciliate nor secreted compounds contributed to the observed loss of infectious HAdV2. Instead, internalization was shown to be the dominant removal mechanism, resulting in the presence of viral particles inside food vacuoles of *T. pyriformis,* as visualized by transmission electron microscopy. The fate of HAdV2 once ingested was scrutinized and no evidence of virus digestion was found over the course of 48 hours. This work shows that *T. pyriformis* can exert a dual role on microbial water quality: while they remove infectious adenovirus from the water column, they can also accumulate and potentially protect infectious viruses from exposure to environmental or engineered stressors.

**Environmental significance:** Human viruses are ubiquitous contaminants of surface water. The fate of human viruses, once discharged into the environment, is modulated by indigenous microorganisms in the surrounding water body. Among microorganisms, the role of protists in controlling virus persistence has been overlooked. Here, we investigate the interactions of the ciliate *Tetrahymena pyriformis* with human adenovirus type 2. We demonstrate that *T. pyriformis* can serve as a sink of adenovirus from the water column, but also act as a reservoir and potential protective barrier for infectious viruses against stressors. *T. pyriformis* ciliates are among the most abundant protists in surface waters. Understanding this dual role of protists is important for assessing and maintaining microbial water quality and infection risks.

## 1. Introduction

Human adenoviruses are frequent contaminants of surface water that can cause diseases such as gastroenteritis, respiratory illness and conjunctivitis^1^. In the aqueous environment, the fate of adenoviruses is in part determined by indigenous microorganisms including protists^2–5^. The interactions of protists with waterborne viruses are manifold, ranging from virus inactivation^6, 7^ to protection and transport^8^, and possibly the induction of higher pathogenicity as demonstrated for a fish virus^9^. For example, the protist *Acanthamoeba polyphaga* was shown to confer protection to human adenovirus type 5 (HAdV5) against chemical disinfection^10^, after internalization of the virions. Conversely, HAdV5 was found to be inactivated by excreted compounds produced by the ciliate *Climacostomum virens*^11^. Depending on the specific interaction, protists may thus serve as a harbor for infectious viruses, or they may contribute to virus control. To fully assess the role of protists on microbial water quality, their interactions with viruses must be better understood.

The type and extent of the interaction depend on both the protist and the virus species^2, 12–14^ highlighting the need to investigate specific protist-virus pairings. Most studies to date have focused on the interactions of waterborne viruses with amoebae^8, 15, 16^ whereas less is known regarding the effects of ciliates. Our recent work has demonstrated that the ciliate *Tetrahymena pyriformis* reduces the infectious titer of different waterborne viruses to differing extents, and that adenoviruses are among the most susceptible toward control by *T. pyriformis*^17^. The mechanisms by which ciliates reduce virus titers, however, are not fully understood, and may include adsorption of viruses to the protist, inactivation by excreted compounds, or ingestion of the virus, with or without subsequent digestion.

Virus ingestion by *T. pyriformis* was demonstrated for vaccinia virus^18^ and rotavirus^19^, whereby the uptake of rotavirus necessitated prior sorption onto the ciliates. For adenovirus, an association of the genome of adenovirus type 3 (HAdV3) with pelleted *T. pyriformis* cells was reported^20^, though it was not determined if the viral particles were absorbed onto, or ingested into the protist cells.

Among viruses taken up by ciliates, bacteriophages were found to be digested and consequently inactivated by *Tetrahymena* sp., possibly serving the ciliates’ nutritional needs^7, 12^. For example, chloroviruses, viruses infecting microalgae, were recently shown to serve as a food source for the ciliates *Halteria sp*. and *Paramecium bursaria*^21^. However, it remains uncertain if ciliates are also capable of digesting and inactivating human viruses including adenoviruses. An early study suggested that influenza virus strains A and B were destroyed upon uptake by *T. pyriformis*^6^. In contrast, the ciliate *Euplotes octocarinatus*^22^ was able to take up human adenovirus type 2 (HAdV2), but the virus exhibited resistance to digestion and infectious virions were still recovered 3 months after ingestion.

Evidently, the removal mechanism and the fate of adenovirus once associated with ciliates remain to be clearly established. In the present study, we provide a comprehensive and mechanistic investigation of the removal of infectious human adenovirus type 2 (HAdV2) by *T. pyriformis*, whereby the term “removal” includes both the inactivation of viruses (i.e. virus decay) and the physical removal of viruses from the water column. *T. pyriformis* was chosen as a model ciliate species since it is present in various water environments, including surface waters^23^ and wastewater^24^. We hypothesized that HAdV2 is removed from water by *T. pyriformis* via ingestion, and that virions are degraded inside its vacuoles, rendering the virus inactive. We conducted experiments to determine the contributions of virus sorption, inactivation by extracellular compounds excreted by protists, ingestion, and digestion to HAdV2 removal from water. Overall, the findings advance our knowledge of the interactions between human viruses and protists, and contribute to a better understanding of the effect of ciliates on microbial water quality.

## 2. Materials and methods

### 2.1 Culturing Tetrahymena pyriformis

The ciliates *Tetrahymena pyriformis (T. pyriformis)* were purchased from the Culture Collection of Algae and Protozoa (CCAP n°1630/1W; Scotland) and were maintained axenically in 75 cm^2^ cell culture flasks (TTP, Milian). The culture medium used was proteose peptone yeast extract (PPYE; 20.0 g × L^−1^ proteose peptone (Bacto peptone, Difco) and 2.5 g × L^−1^ yeast extract (BioChemika) in Mili-Q water, autoclaved and stored at 4°C). The cultures were maintained in a 28°C incubator to promote ciliate growth^25^. Every week subcultures were prepared by re-spiking 1 ml of the previous culture into 19 ml of fresh PPYE.

### 2.2 Virus propagation, purification and infectivity quantification

Human mastadenovirus C type 2 (HAdV2, kindly provided by Rosina Girones, University of Barcelona) was propagated by infecting sub-confluent monolayers of A549 cells (Rosina Girones, University of Barcelona) in 150 cm^2^ cell culture flasks (TTP, Milian). A549 cells were cultured with Dulbecco’s Modified Eagle’s medium (DMEM) containing 10% Fetal Bovine Serum (Gibco, Frederick) and 1% Penicillin/Streptomycin (Gibco, Frederick), and maintained in an incubator at 37°C and 5% CO_2_. Once infected, cells underwent three freeze-thaw cycles (−80°C/25°C) to release virions. The resulting solution was centrifuged for 15 minutes at 1’700 x *g* to remove cell debris. The supernatant was filtered through a 0.22 µm syringe filter (Millex-GV, 0.22 µm, PVDF, 33 mm, Gamma-Sterilized) and the filtrate was washed with phosphate-buffered solution (PBS; 0.71 g × L^−1^ of Na_2_HPO_4_, 0.58 g × L^−1^ of NaCl in deionized water, followed by autoclave, pH 7.4; Acros organics) three times using 100 kDa amicon filters (Merck, Millipore) and centrifuging at 3’000 × *g* for 6 min. The resulting HAdV2 stock solution was maintained at −20°C until use. Infectious HAdV2 concentrations were enumerated by the most probable number (MPN) assay as described elsewhere^26^ and were expressed as most probable number of cytopathic units per milliliter (MPNCU × ml^−1^). The Rstudio software (version 1.0.153)^27^ was used to determine the infectious virus titers from the MPN assay. The limit of quantification (LOQ) of the MPN assay, defined as the concentration that results in one cytopathic unit in the undiluted sample, and none in all subsequent dilutions of the MPN assay^28^, corresponded to 15 MPNCU × ml^−1^.

### 2.3 Viral DNA quantification

Total (infectious and inactivated) HAdV2 concentrations were quantified by quantitative polymerase chain reaction (qPCR) and were expressed in genomic copies per milliliter (GC × ml^−1^). Extractions of viral nucleic acids from 140 µl samples were performed with the QIAamp Viral RNA Mini Kit (Qiagen), according to the manufacturer’s protocol. The resulting nucleic acids were eluted in 60 µl of AVE buffer and stored at −20°C prior to analysis by qPCR. A negative extraction control using ultra-pure distilled water (Invitrogen) was included in each extraction batch and always yielded negative results. qPCR reactions were performed using the TaqMan Environmental PCR Master Mix (Applied Biosystems, Part n° 4396838) with an amplicon size of 81 base pairs and the following forward primer, reverse primer and probes^29^: F:5’CWTACATGCACATCKCSGG-3’; R:5’-CRCGGGCRAAYTGCACCAG-3’; 5’FAMCCGGGCTCAGGTACTCCGAGGCGTCCT-BHQ1-3’.

The volume of extracted nucleic acid was 10 µl and the final reaction volume was 25 µl per sample. Thermocycling reactions were performed in Mic Real-Time PCR devices (Bio Molecular Systems). Each reaction was performed in technical duplicate. Each qPCR run contained a no-template control consisting of ultra-pure distilled water, which yielded negative results. Standard curves were prepared by 10-fold serial dilutions of gblock gene fragments (Integrated DNA Technologies) in TE buffer (Invitrogen), ranging from 10 to 1 × 10^7^ GC × µl^−1^ and were included in each run. The thermocycler program consisted of 10 min at 95°C followed by 40 cycles (15 s at 95°C and 1 min at 60°C). The micPCR software (version 2.10.0) was used to acquire Cq values and for absolute quantification. Pooled standard curves with several replicates of low concentration standards were analyzed in R using the Generic qPCR Limit of Detection (LOD) / Limit of Quantification (LOQ) calculator^30^. The LOQ of the qPCR assay, defined as the lowest standard concentration with a coefficient of variation ≤ 35%, was 287 genomic copies/reaction. The standard curve used in the sorption experiments exhibited a slope of −3.377, a PCR efficiency of 98 % and an R^2^ of 0.9988. No evidence of PCR inhibition in the samples was found, as determined by qPCR analysis of both undiluted and ten-fold diluted samples.

### 2.4 Co-incubation experiments

The effect of *T. pyriformis* on HAdV2 infectivity was determined in co-incubated experiments over the course of 48 h or 72 h. Prior to co-incubation, *T. pyriformis* were washed and starved. Washing was performed following the protocol of Pinheiro et al.^31^ except that the rinsing solution was sterilized moderately hard synthetic freshwater^32^ (MHSFW; 96.0 mg × L^−1^ NaHCO_3_ (Acros organics), 60.0 mg × L^−1^ CaSO_4_*2H_2_O (prepared with chemical grade reagents), 60.0 mg L^−1^ × MgSO_4_ (AppliChem) and 4 mg × L^−1^ KCl (AppliChem) in deionized water). After two washes, ciliates were resuspended in MHSFW and incubated at room temperature to induce starvation. Starvation time ranged from 15 h to 21 h, unless mentioned otherwise. Ciliate concentrations were measured by taking the average of three counts of 10 µl *T. pyriformis* samples using a Neubauer counting chamber (Bright-line, Hausser-scientific) and an inverted microscope (Olympus CK X41). Only the number of actively moving ciliates, therefore non-encysted ciliates, were counted. Co-incubation experiments were carried out in 50 ml plastic tubes (Falcon, Greiner) containing 3 to 10 ml of *T. pyriformis* solution in MHSFW at an average concentration ranging from 1 × 10^4^ to 1 × 10^5^ cells × ml^−1^. A 100 µl aliquot of HAdV2 virus stock was spiked into each reactor to reach an initial virus concentration between 4 × 10^5^ MPN × ml^−1^ and 2 × 10^5^ MPN × ml^−1^. The resulting virus-to-protist ratio ranged between 4:1 and 30:1. Control reactors consisted of 3 to 10 ml of MHSFW spiked with 100 µl of the virus but without *T. pyriformis*. Throughout the co-incubation experiments, aliquots of 200 µl were taken at time points 0, 24 h, 48 h or 72 h, were filtered through 0.22 µm syringe filter to remove protists, and were quantified for virus infectivity by the MPN assay. This filtration step was performed to avoid confounding effects of protists during quantification. The 0.22 µm filtrate represents the viral fraction and control experiments confirmed that the majority of viruses were present in this filtrate. Each experiment was carried out in triplicate.

Net virus removal (or net virus loss) by *T. pyriformis* was quantified on a log_10_ scale as the loss in the infectious virus concentration (log_10_ (C/C_0_)) over 48 hours, unless mentioned otherwise. Alternatively, the loss in genomic copies (log_10_ (N/N_0_)) was measured if infectivity measurements were not experimentally accessible.

The net removal was calculated as the virus loss in the experiment (log_10_ (C_exp_/C_exp,0_)) corrected for the virus loss measured in the *T. pyriformis*-free control (log_10_ (C_cont_/C_cont,0_)), (Equation 1).

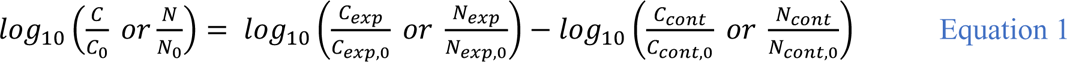

### 2.5 Virus inactivation by excreted compounds

To investigate if ciliates were excreting compounds into the solution that would inactivate HAdV2 (i.e. inactivating compounds), *T. pyriformis* cells were first washed and starved as described above, resulting in 20 ml of starved cells in MHSFW at a concentration of 8 × 10^4^ cells × ml^−1^. After 21 hours, 10 ml of the solution were filtered through a 0.22 μm syringe filter to remove ciliates. The obtained filtrate mimicked solution conditions prior to virus exposure in the default co-incubation experiment. 3 ml of filtrate were amended with 100 µl of HAdV2, resulting in an initial virus concentration of 3 × 10^6^ MPN × ml^−1^. Controls consisted of unfiltered solutions containing live *T. pyriformis*. Aliquots of 200 μl were taken at 0 h, 48 h and 72 h and were filtered through a 0.22 μm syringe filter. HAdV2 infectivity was measured by MPN assay.

### 2.6 Virus sorption onto *T. pyriformis*

To assess the extent of virus adsorption onto the ciliate, *T. pyriformis* were killed by adding 1 mL of 10% neutral buffered formalin (Sigma-Aldrich) to 3 ml of a solution containing 8.5 × 10^4^ ciliates × ml^−1^ in MHSFW, followed by incubation at 25°C for 10 min. Because the experimental concentration of formalin caused detrimental effects on A549 cells, samples were diluted at least 100-fold prior to enumeration by MPN assay. Controls consisted of live *T. pyriformis* in 4 ml of MHSFW. To each solution, 100 µl of HAdV2 from a stock at 1.7 × 10^8^ MPN × ml^−1^ were added, resulting in an initial virus concentration of 2 × 10^6^ MPN × ml^−1^. In samples containing formalin, the initial virus concentration was immediately reduced to 10^3^ MPN × ml^−1^ by the addition of formalin. After this initial drop due to chemical action, the physical sorption could be monitored. Aliquots of 300 μl were taken at times 0 h and 48 h and both HAdV2 infectivity and genomic copy numbers and were measured.

### 2.7 Cytochalasin B treatment

The contribution of virus ingestion to the observed virus removal by *T. pyriformis* was assessed in experiments containing an actin filament formation inhibitor^33^, Cytochalasin B (Cyt B; produced by *Drechslera dematioidea*, Sigma-Aldrich). To ingest particles, ciliates usually perform phagocytosis on particles > 250-500 µm^31, 34^ while they use pinocytosis to acquire smaller particles and nutrients from their surrounding fluid, before they end up in membrane vesicles or food vacuoles^31, 35^. Cyt B can inhibit both phagocytosis and pinocytosis^29, 33^. Experiments were conducted using 4 to 5 ml of *T. pyriformis* in MHSFW, which were washed as described above and starved for 48 h. Cyt B addition to the samples was achieved following a protocol published eslewhere^31^, with minor modifications. Specifically, 1 mg of Cyt B was dissolved into 100 μl of dimethyl sulfoxide (DMSO; Sigma-Aldrich) and 15 μl aliquots of the resulting Cyt B solution were added to each reactor. Finally, the reactor was amended with 100 μl of HAdV2. Initial protist concentrations ranged between 4.8 × 10^4^ cells × ml^−1^ and 1.4 × 10^5^ cells × ml^−1^ and initial virus concentrations ranged between 9.8 × 10^5^ MPN × ml^−1^ and 3.6 × 10^6^ MPN × ml^−1^. Because the effect of Cyt B is time-restricted^33^, the experimental solutions were re-spiked with 22.5 μl of Cyt B solution after 15 h of co-incubation with the virus. Aliquots of 350 μl were taken at times 0 h and 21 h and were filtered through a 0.22 μm syringe filter before quantification by MPN assay. The Cyt B did not influence the downstream virus enumeration nor protist growth. Controls consisted of Cyt B-free solutions and *T. pyriformis*-free solutions.

To confirm that Cyt B led to a reduction in food vacuoles, 500 µl of *T. pyriformis* were treated with either 2.5 µl of Cyt B/DMSO solution or with 2.5 µl DMSO alone. After 10 minutes of incubation at room temperature, 2 ml of 1% carbon ink (Aqua ink Graph, Jumbo) were added to each sample to stain the food vacuoles. Sample aliquots (40 µl) were taken at regular time points over 40 h and fixed with 10 µl of 1% glutaraldehyde each (Electron Microscopy Sciences). The number of food vacuoles in 10 cells was counted under light microscopy. Experiments were conducted in duplicate.

### 2.8 Quantification of egestion of HAdV2 and of infectious HAdV2 titer inside *T. pyriformis*

The infectivity of HAdV2 taken up by *T. pyriformis* was quantified to determine if viruses can be digested by protists. Specifically, eight reactors (50 mL) containing 5 ml of overnight starved ciliates at a concentration of 1.6 × 10^5^ cells × ml^−1^ in MHSFW were prepared. Each reactor was amended with HAdV2 to a final concentration of 6.8 x 10^4^ MPN × ml^−1^ and were incubated at room temperature for one hour. Then, each solution was centrifuged at 400 × *g* for 10 minutes. The resulting cell pellets were washed twice with 5 ml of MHSFW by centrifuging at 400 × *g* for 6 min and discarding the supernatant each time. This step allowed to remove extracellular virions (though extracellular vesicles could also be washed out). Each wet pellet (0.5 ml) was then transferred to a new reactor containing 4.5 ml of virus-free MHSFW, to halt further virus uptake. The protist concentration after transfer was around 10^5^ cells × ml^−1^ and remained stable for the duration of the experiment. At different times following transfer (0 h; 12 h; 24 h and 48 h), aliquots of 100 µl were sampled to determine the concentration of egested virus in solution by MPN. To enumerate the ingested viruses, the remaining reactor content was pelleted by centrifuging at 400 × *g*. The supernatant was removed from the pellet, and the residual aqueous volume was recorded for each sample, to determine the number of infectious viruses present in residual supernatant surrounding the pellet. The pellet was disrupted by three cycles of −20°C / 37°C freezing/heating. Of the resulting suspension, 100 µl were sampled and the infectious HAdV2 concentration was measured. Finally, the infectious HAdV2 number in the pellet suspension was then corrected for the number of infectious virus resulting from the residual supernatant. Figure S1† summarizes the procedure for this experiment.

To examine if (infectious) virions could be released from protists after longer co-incubation times, we exposed washed and starved *T. pyriformis* (4 × 10^5^ cells × ml^−1^) to HAdV2 (1.6 × 10^6^ MPN × ml^−1^) for 72 h instead of 1 h. Protists were then rinsed and transferred into 3 ml of virus-free MHSFW. Aliquots of 350 µl were sampled over the course of 24 h and were analyzed for viral genomic copies and infectivity.

### 2.9 Transmission electron microscopy and tomography

For visual confirmation of internalized virus particles, starved *T. pyriformis* were co-incubated with HAdV2 in MHSFW for 50 minutes at room temperature and at two different virus-to-protist ratios (Table S1†). Viruses were spiked into 3 ml or 5 ml of *T. pyriformis* solutions at the beginning of the co-incubation, and again after 30 minutes. Control experiments without viruses were also conducted. Protist suspensions were then fixed in solutions of 2.5 % glutaraldehyde (Electron Microscopy Sciences) in phosphate buffer (0.1 M, pH7.4) (Sigma) for 1 hour at room temperature and were centrifuged at 755 x *g* for 2 minutes. The resulting pellets were directly postfixed by a fresh mixture of osmium tetroxide 1% (Electron Microscopy Sciences) with 1.5% of potassium ferrocyanide (Sigma) in phosphate buffer for 1 hour at room temperature. The samples were then washed three times in distilled water and spun down in low melting agarose (2% in H_2_O; Sigma), let to solidify on ice, cut into 1 mm^3^ cube and dehydrated in acetone solution (Sigma) at graded concentrations (30%-40 min; 70%-40 min; 100%-2 × 1 h). This was followed by infiltration in Epon (Sigma) at graded concentrations (Epon 1/3 acetone-2 h; Epon 3/1 acetone-2h, Epon 1/1-4 h; Epon 1/1-12 h) and finally polymerized for 48 h at 60°C in oven. Semi-thin sections of 250 nm were cut on a Leica Ultracut (Leica Mikrosysteme GmbH) and picked up on a 2 × 1 mm copper slot grid (Electron Microscopy Sciences) coated with a polystyrene film (Sigma). Gold particles of 10 nm in diameter were deposited on both faces of the grid for tomogram alignment and the sections were post-stained with 2% uranyl acetate (Sigma) in H_2_O during 10 minutes. Finally, the grids were rinsed several times with H_2_O and then with Reynolds lead citrate in H_2_O (Sigma) for 10 minutes, followed by several additional rinses with H_2_O.

We identified HAdV2 particles based on their icosahedral structure, their size, and example TEM images of HAdV2 particles in infected cells reported in the literature^36^. In addition, we thoroughly investigated the food vacuoles in cells that were not exposed to HAdV2 (negative control) to exclude any viral particles similar to HAdV2 in shape that could already be commensal to the protists.

Dual-axis tomograms were collected on a JEOL 2100Plus electron microscope using a Fischione dual-axis tomography holder Model 2040 (Fischione Instruments) and a TVIPS XF416 camera (TVIPS GmbH) using SerialEM software^37^ at an accelerating voltage of 200kV, and 30000X of magnification (pixel size of 0.77 nm). Dual-axis tomograms reconstruction was performed using etomo with SIRT projection (4 iterations) from the IMOD software^38^.

### 2.10 Data analysis

Statistical analyses were performed GraphPad Prism 9^39^ version 9.0.2. and results were considered significant for p < 0.05.

## 3. Results

### 3.1 Infectious HAdV2 particles in aqueous solution are efficiently reduced by *T. pyriformis*

HAdV2 removal was quantified either in co-incubation with *T. pyriformis* in MHSFW or in MHSFW alone, as described in section 2.4. In the presence of *T. pyriformis,* we observed a drop in the infectious titer of HAdV2, as compared to the ciliate-free control (Figure 1A). Consequently, in the next experiments we mainly used a co-incubation time of two days since this was sufficient to detect a reduction greater than 99% of the initial infectious virus load In three independent experiments with a virus-to-protist ratio ranging from 4:1 to 30:1, the net effect of *T. pyriformis* on HAdV2 was always significant (Figure 1B; see Table S2† for detailed statistical results). However, we observed slight variability in the magnitude of the mean effect which ranged between 2.6 log_10_ and 3.3 log_10_ over 48 h. The net average removal was 2.8 log_10_ over 48 h. The virus-to-protist ratio varies less than 1-log order of magnitude. Based on our previous study^17^ such a ratio change has no significant impact on the virus removal rate.

**Figure 1:**
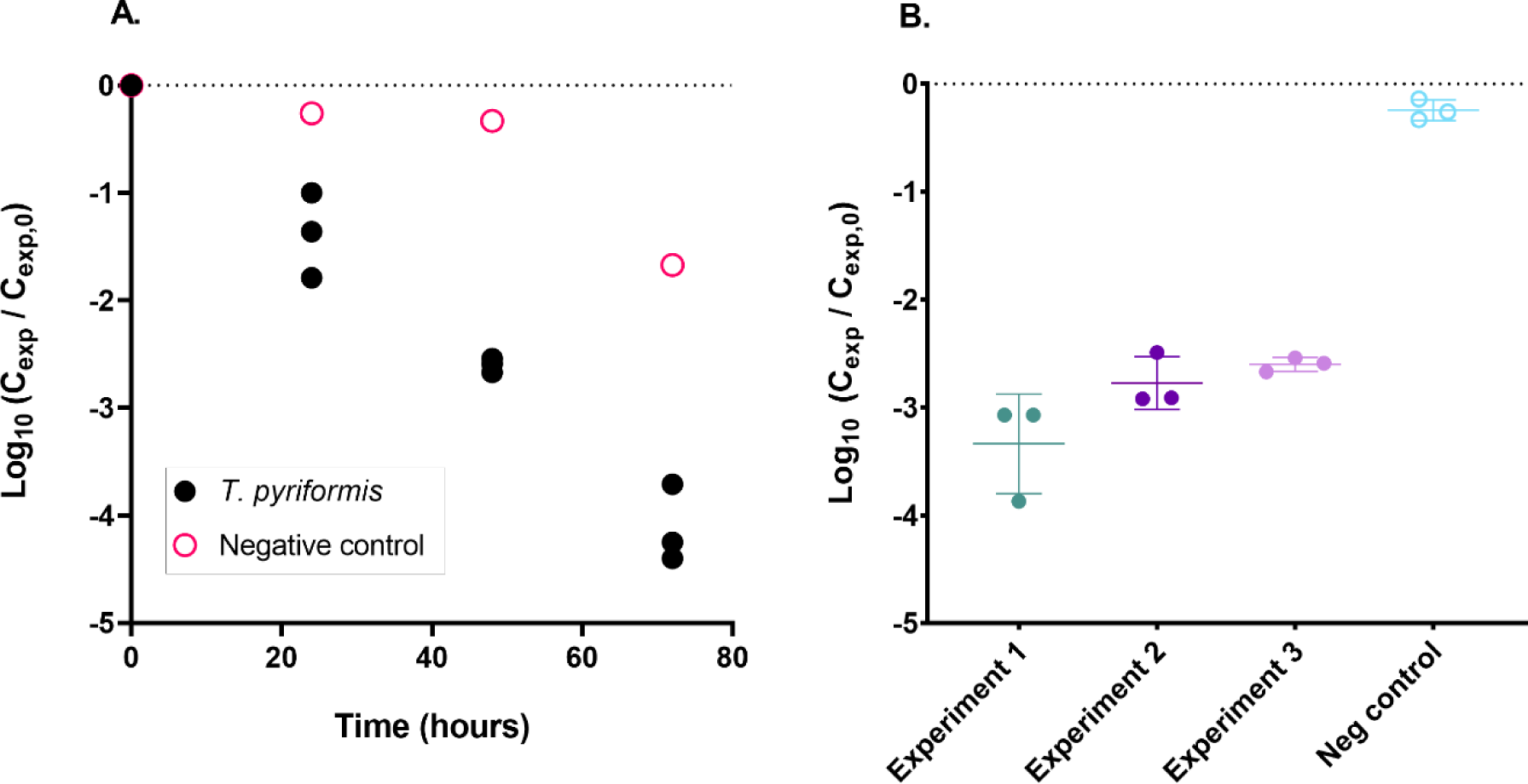
Monitoring of HAdV2 removal by *T. pyriformis*. **(A)** Log_10_ reduction in HAdV2 infectivity during co-incubation in MHSFW containing *T. pyriformis* or in MHSFW alone. The initial virus titer was 4 × 10^5^ MPN × ml^−1^ and the initial *T. pyriformis* concentration was 1 × 10^5^ cells × ml^−1^. **(B)** Log_10_ reduction in HAdV2 infectivity after co-incubating *T. pyriformis* in MHSFW with HAdV2 for 48 h. Three experiments were carried out with different virus-to-protist ratios (30:1 for experiment 1 and 4:1 for experiments 2 and 3). Data for experiment 3 correspond to data shown in Figure 1A. The horizontal bars indicate the mean log_10_ reduction of three experimental replicates and the associated standard deviation. See Table S2† for results of the ANOVA analysis to compare reduction in infectivity in the different experiments.

Yet, to minimize experimental variability, all analyses hereafter were made for replicate experiments carried out with the same protist subculture (as defined in section 2.1) and the same virus-to-protist ratio.

### 3.2 Contribution of excreted compounds, sorption and ingestion to virus removal by *T. pyriformis*

When co-incubating HAdV2 with *T. pyriformis* filtrate, retaining only potential excreted compounds as described in section 2.5, the HAdV2 infectious titer remained stable over 48 h (Figure 2; unpaired two-tailed t-test: t(13.33) = 4.897; p < 0.0001). In contrast, the removal measured in samples containing *T. pyriformis* was ≥ 3 log_10_ during this time period. Thus, the presence of ciliates is required to achieve HAdV2 removal.

**Figure 2:**
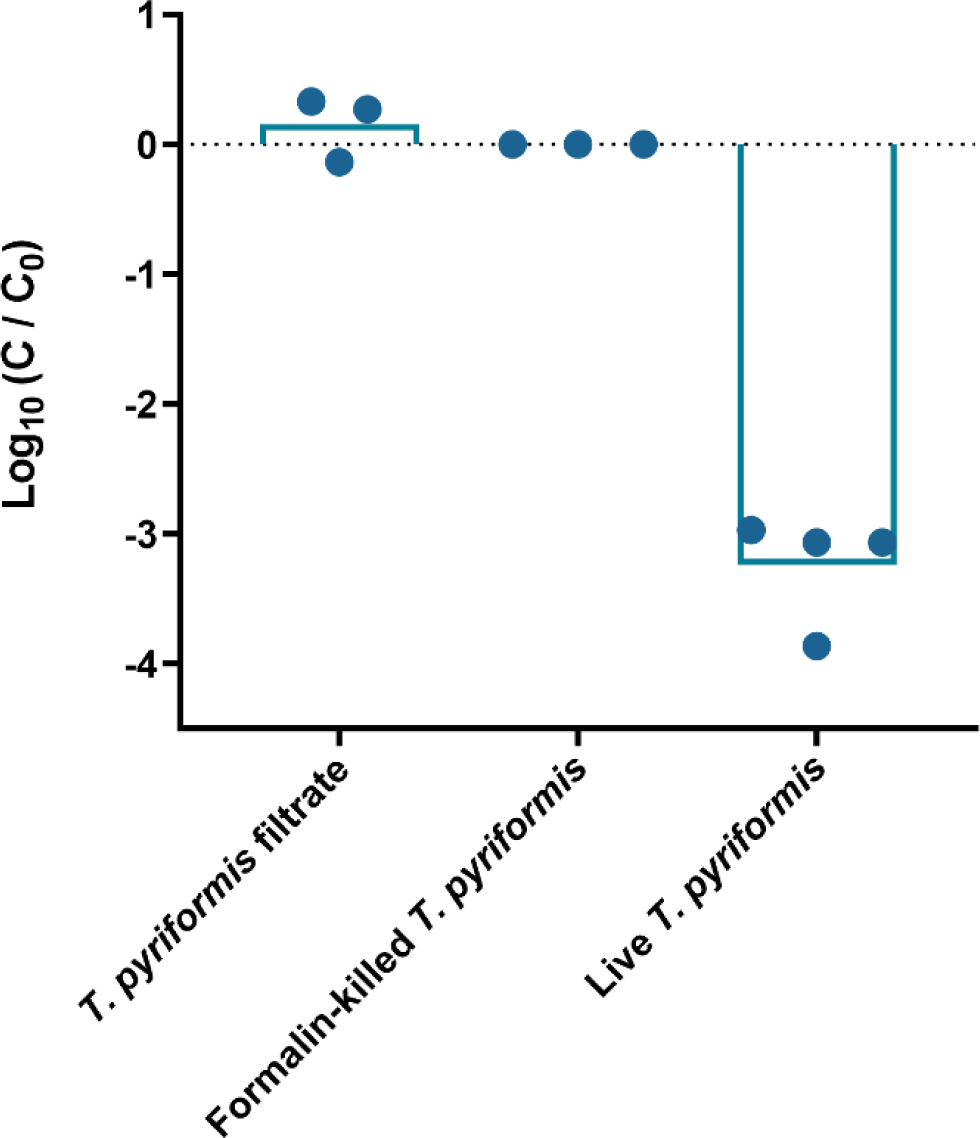
Effect of secreted compounds and sorption on HAdV2. Net log_10_ decrease of HAdV2 infectivity when formalin-killed *T. pyriformis*, the filtrate of the starving *T. pyriformis* cultures containing only excreted compounds, or living *T. pyriformis* were co-incubated with HAdV2 for 48 h. The initial virus titer was 2 × 10^6^ MPN × ml^−1^ and the initial *T. pyriformis* concentration was 8 × 10^4^ cells × ml^−1^. The vertical bars indicate the mean log_10_ reduction of three or more replicates.

Sorption tests revealed that after 48 h of co-incubation of formalin-killed *T. pyriformis* with HAdV2, no infectious virus removal occurred, while a 3 log_10_ decrease in infectivity was measured in reactors containing living *T. pyriformis* (Figure 2; unpaired two-tailed t-test: t(13.13) = 5; p < 0.0001). Likewise, the genome copy number, representing the total (infectious plus inactivated) number of viral particles, remained stable in formalin-killed *T. pyriformis*, but underwent a net removal of 1.8 log_10_ in the presence of live *T. pyriformis* (Table S3†). This suggests that neither infectious nor inactivated virus was adsorbed onto *T. pyriformis*.

As a final mechanism, we investigated the role of virus ingestion in HAdV2 removal. We first determined that the feeding inhibitor Cyt B was efficient in reducing the production of vacuoles formed by *T. pyriformis*. After three hours, the number of vacuoles per ciliate exposed to Cyt B was seven-fold less than in the untreated organisms (Figure 3A). Visual confirmation of food vacuole impairment by light microscopy can be found in supplementary data (Figure S2†).

**Figure 3:**
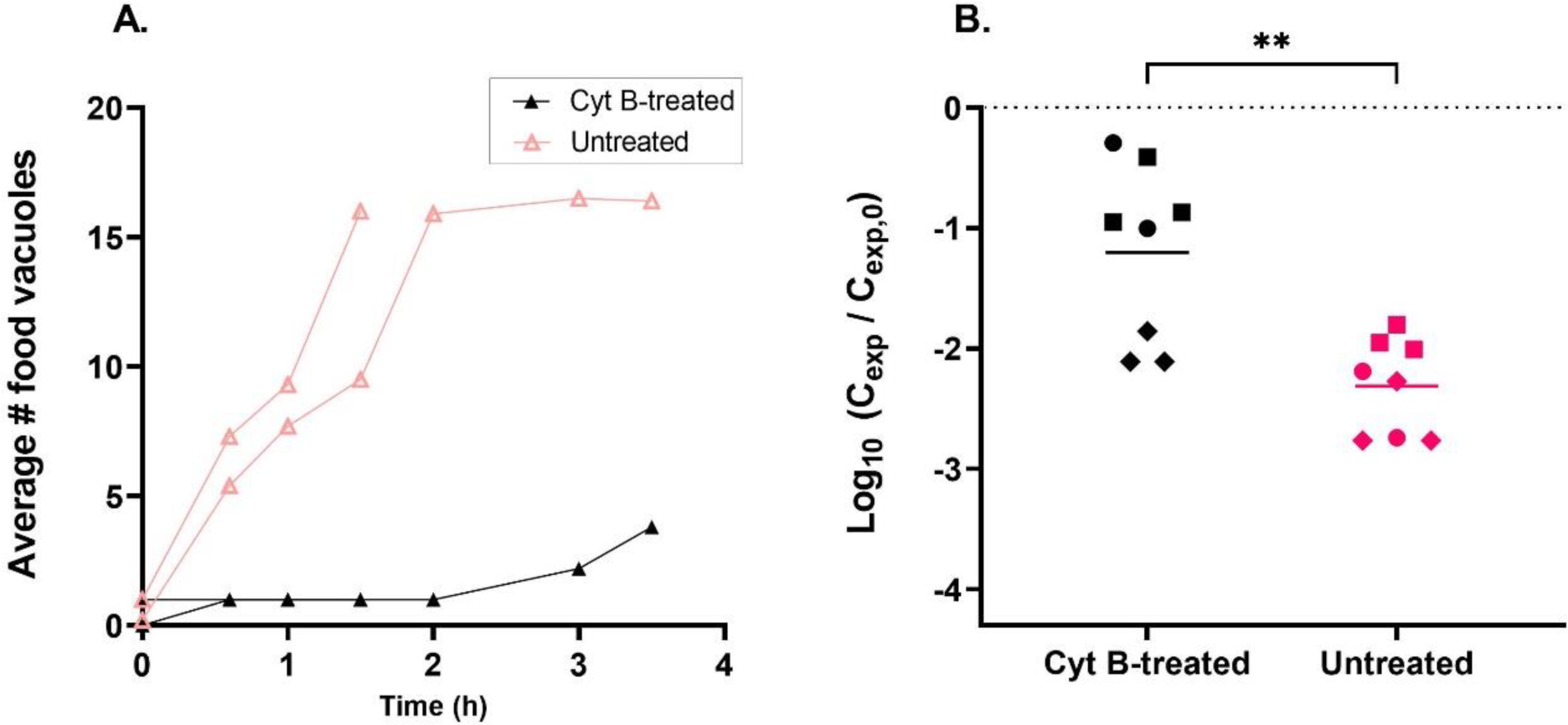
Effect of Cytochalasin B on food vacuole formation and removal of HAdV2. **(A)** Monitoring of food vacuole formation in Cyt B-treated and untreated *T. pyriformis* over a 4-hour time course (in duplicate). Cyt B was re-spiked over the course of an experiment as detailed in section 2.7. Visual confirmation by light microscopy of the action of Cyt B is found in Figure S2† **(B)** Log_10_ reduction in HAdV2 infectivity after co-incubation with Cyt B-treated or untreated *T. pyriformis* over 21 h in MHSFW. Each symbol corresponds to an independent experiment carried out in triplicate. The horizontal bars represent the mean of all replicates. The asterisks indicate a p-value < 0.01 (unpaired two-sided t-test with a validated assumption of equal variances).

Correspondingly, when HAdV2 was incubated with Cyt B-treated protists, the reduction in virus infectivity was significantly lower than in samples with untreated *T. pyriformis* (unpaired t-test, two-tailed: t(14) = 3.782; p-value = 0.0020) (Figure 3B). These results indicate that internalization by phagocytosis, pinocytosis, or both plays a role in the removal of infectious virions by *T. pyriformis*.

To obtain further confirmation of ingestion, we analyzed cell compartments of *T. pyriformis* with and without co-incubation (1 h) with HAdV2 by transmission electron microscopy. We detected virions in several food vacuoles of *T. pyriformis* that had been co-incubated with HAdV2 (Figure 4 and Figure S3†). We observed up to 13 viral particles per slice of food vacuole, as illustrated in Figure 4A. In contrast, no viruses were found in *T. pyriformis* from the virus-free control experiments. Images of additional vacuoles are shown in Figure S3†. These findings were further confirmed by analyzing slices of a given food vacuole, in which we could detect several viral particles at different depths. The reported observations are available as tomograms under https://doi.org/10.5281/zenodo.6808144, †.

**Figure 4:**
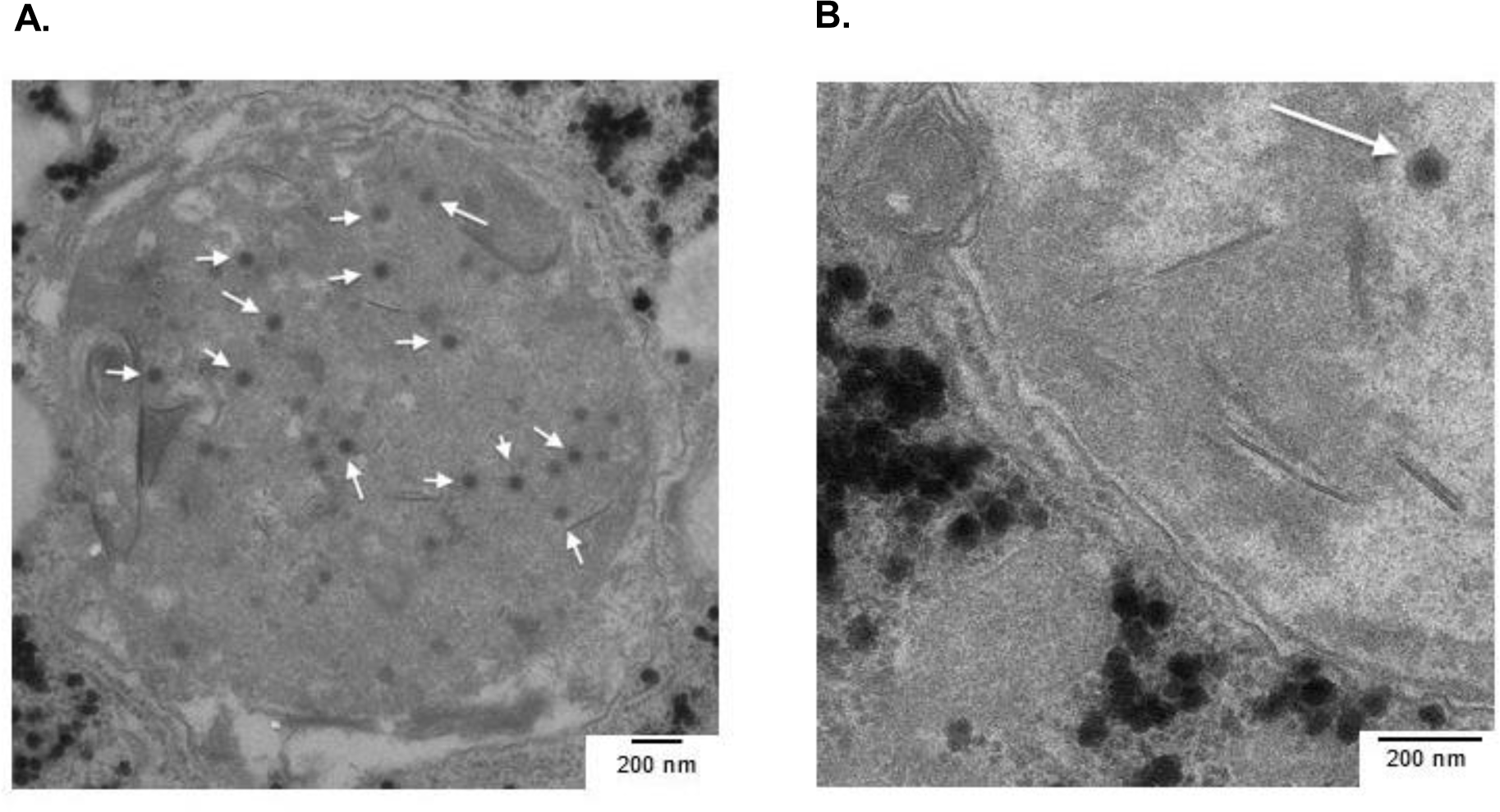
Transmission electron microscopy images of HAdV2 particles inside a *T. pyriformis* food vacuole. **(A)** Example of a food vacuole of *T. pyriformis* co-incubated with HAdV2 for 50 minutes at a virus-to-protist ratio of 3’000: 1. Examples of virions are indicated with white arrows; **(B)** Enlarged image of a single HAdV2 particle inside a *T. pyriformis* food vacuole.

### 3.3 Assessment of HAdV2 fate upon ingestion by *T. pyriformis*

To measure virus egestion and digestion, HAdV2 was first co-incubated with *T. pyriformis* for 1 h. As shown above, this co-incubation period was sufficient to accumulate observable numbers of HAdV2 virions within *T. pyriformis* food vacuoles (Figure 4). Upon transfer and resuspension of pelleted *T. pyriformis* in fresh MHSFW (see Figure S1†), approximately 10^4^ MPN × ml^−1^ infectious HAdV2 were measured in the solution, likely stemming from residual experimental solution transferred with the pellet. This virus concentration remained stable over the course of 48 h (Figure 5A), indicating that there was no measurable net egestion of infectious HAdV2 from *T. pyriformis.* Similarly, the use of a longer initial co-incubation time (72 h instead of 1 hour) resulted in neither measurable egestion of infectious nor total HAdV2 virions (data not shown).

**Figure 5:**
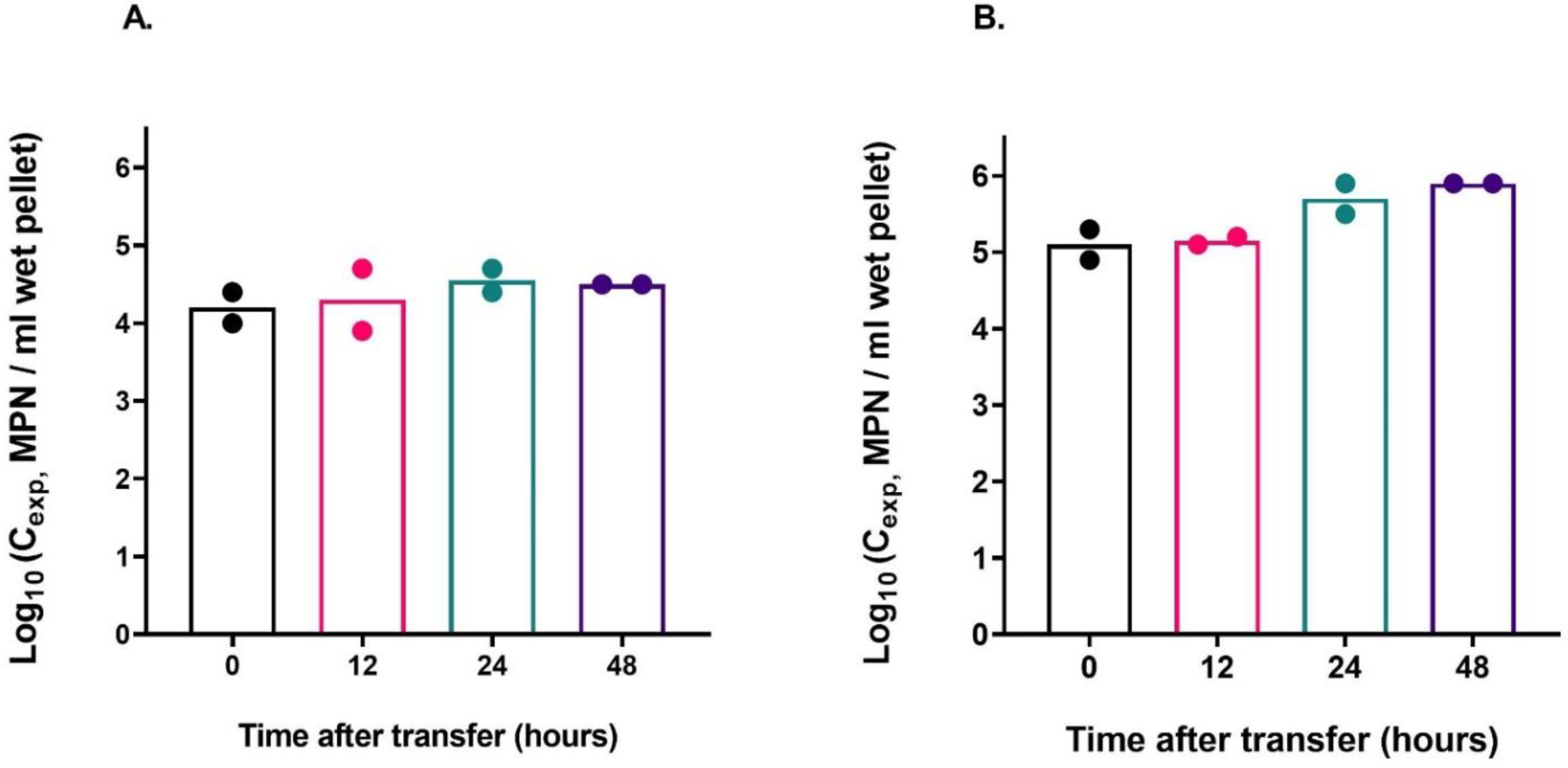
Monitoring of virus egestion (A) and digestion (B) over time following the method described in Figure S1†. **(A)** Log_10_ concentration of infectious HAdV2 measured in solution of experiments in which *T. pyriformis* were co-incubated with HAdV2 for 1 hour before transfer into fresh MHFSW. **(B)** Log_10_ concentration of infectious HAdV2 measured in the lysed pellets at different times after 1 hour-co-incubation of *T. pyriformis* with HAdV2 and transfer to fresh MHFSW.

Consistent with a lack of net egestion, a high concentration of infectious HAdV2 was retained within *T. pyriformis* (Figure 5B). Infectious HAdV2 concentrations in the pelleted ciliates remained > 10^5^ MPN × ml^−1^ of wet pellet over 48 h after transfer into fresh MHSFW. No statistically significant difference was observed between the virus concentration recovered at different times post-transfer (one-way ANOVA with Tukey’s multiple comparisons’ tests; p-values > 0.05; see full statistical results in Table S4†).

## 4. Discussion

### 4.1 Mechanism of HAdV2 removal by *T. pyriformis*

Human adenoviruses encounter ciliates in various aqueous environments; to assess the role of ciliates in modulating microbial water quality, it is critical to understand how their presence affects adenovirus infectivity and circulation. In the present work, the results of HAdV2 and *T. pyriformis* co-incubation experiments indicated that HAdV2 viral particles could efficiently be removed from water via *T. pyriformis*. The net reduction in infectious virus by 1 × 10^4^ to 1 × 10^5^ ciliate cells × ml^−1^ was on average 2.8 log_10_ within two days (Figure 1).

Neither physical sorption of the viral particles onto the protist cells, nor secretion of inactivating compounds was found to contribute to virus removal under the experimental conditions used herein (Figure 2). We cannot exclude that the lack of sorption onto *T. pyriformis* resulted from a change in surface adsorption properties due to formalin treatment. However, virus removal was also reduced if the production of food vacuoles was inhibited, which confirms that sorption is not the dominant removal mechanism.

The lack of inactivation induced by excreted compounds is consistent with findings for vaccinia virus^18^ when co-incubated with *T. pyriformis.* However, our results are contrary to previous suggestions that a chemical produced by ciliates contributes to the inactivation of human adenovirus^11^. It thus remains to be tested if protists excrete inactivating compounds under different solution conditions than the ones employed herein, or if other adenovirus species are more sensitive to those compounds.

Instead, virus removal could be attributed to ingestion of viruses by *T. pyriformis*. Two lines of evidence support this finding. First, if phagocytosis and pinocytosis were inhibited by treatment of *T. pyriformis* with Cyt B, the virus removal by the ciliate was significantly reduced (Figure 3B). And second, HAdV2 particles could be visualized inside food vacuoles of co-incubated protists (Figure 4 and Figure S3†). We thus confirm our hypothesis that the main mechanism leading to HAdV2 removal by *T. pyriformis* is internalization. This conclusion is consistent with the suggestion of Sepp et al.^20^ that HAdV3 viral particles could be taken up and located in food vacuoles of *T. pyriformis*.

Cyt B effectively impaired actin filament formation and reduced vacuole number formation (Figure 3A, Figure S2†). It is an appropriate chemical to inhibit protistan feeding^33^ and has proven reliable to study grazing on several bacteriophages^7,31^. This treatment, however, does not allow to disentangle the different forms of internalization. Further studies should address the point of entrance of HAdV2 virions (i.e. cytostome or membrane surface). Yet, considering the diameter of HAdV2 particles (90 nm), pinocytosis is likely to be the dominant internalization mechanism of HAdV2. Pinocytosis was also suggested as a removal mechanism of T4 bacteriophage by *Tetrahymena thermophila*^31^. Incidentally, for other protist species, virions have been shown to accumulate in cell compartments other than food vacuoles, as recently demonstrated for reovirus, which was detected in the nucleus of amoeba^16^. This confirms that alternative internalization routes to pinocytosis exist.

### 4.2 Fate of the virus following ingestion

We found no evidence of net egestion of (infectious) viral particles from *T. pyriformis* after co-incubation with HAdV2 (though we cannot rule out constant exchange of virus between ciliates and the media). Instead, once ingested, the virus particles are accumulated rather than released by *T. pyriformis* (Figure 5A). These data corroborate the study of Sepp et al.^20^, who co-incubated *T. pyriformis* with HAdV3 and then successively passaged the protists in virus-free medium. Authors could not detect viral antigens after the second passage, consistent with limited virus egestion.

In addition, our findings showed no reduction in the infectious concentration of internalized viruses over 48 hours (Figure 5B), indicating no digestion of HAdV2. Our hypothesis of virus inactivation upon internalization is thus refuted, at least on the time scale considered. Our data are in agreement with a study showing that vaccinia virus could persist inside *T. pyriformis* for 10 days^18^, but are discrepant from reports of ingested influenza virus inactivation^6, 40^ and recent reports of ingested chloroviruses by other ciliates^21^.

Virus digestion by protists is known to be both virus- and protist-specific, as shown for bacteriophages^12, 13^. Given that food vacuoles are acidic upon ingestion^41^, a virus’ propensity to become digested may depend on its sensitivity to low pH. Alternatively, the discrepancies in virus fate reported in different studies may also be a result of experimental artifacts in conjunction with the pellet-washing protocols used. In the present study, we found that washing the pelleted protists was a critical step in assessing the number of infectious viruses internalized, since cells could lyse during washing and centrifugation, thereby releasing viral particles.

### 4.3 Implications for pathogen control in aquatic environment

While *T. pyriformis* can act as a biofilter for HAdV2 removal in MHSFW (Figure 1), its ability to store infectious viruses (Figure 5) could become a liability during water and wastewater treatment. First, *T. pyriformis* may serve as a reservoir of infectious virions, which – upon disruption – may cause the release of locally high concentrations of infectious virions. Second*, T. pyriformis* could protect infectious HAdV2 from water disinfectants or environmental stressors. While such protective effects were not investigated in this work, previous work has reported that bacteriophages internalized by ciliates could be shielded from UV disinfection^42^, and that adenovirus type 5 ingested by *Acanthamoeba polyphaga* were protected from chlorine disinfection^10^. And finally, *T. pyriformis* could aid the transport of infectious HAdV2 virions throughout a water body, or catalyze the transfer of infectious viruses through aquatic food chains, as previously reported for bacteriophages and enterovirus^43^.

## 5. Conclusions

This work combines a series of experiments to decipher the mechanisms underlying the fate of HAdV2 in presence of the model ciliate *T. pyriformis.* We determined that:

(i) *T. pyriformis* efficiently removed infectious HAdV2 from aqueous solution.
(ii) the main responsible mechanism for HAdV2 removal was internalization.
(iii) neither compounds excreted by *T. pyriformis* nor sorption onto the ciliates affected infectious HAdV2 concentrations.
(iv) ingested viruses were located inside *T. pyriformis* food vacuoles.
(v) internalized HAdV2 could be recovered from *T. pyriformis* in an infectious state.

These findings have implications for water quality. *T. pyriformis* serve a dual role as a modulator of microbial water quality. On the one hand it removes infectious HAdV2 from the water column. On the other hand, it acts as a reservoir for infections viruses. Future work should investigate if internalization by *T. pyriformis* leads to protection from natural (e.g., sunlight) or engineered (e.g., disinfectants) stressors, or if water treatment processes can lead to a rupture of *T. pyriformis* cells and subsequent release of infectious viruses in the environment. The results of this work also have implications for surface water ecology. The fact that HAdV2 remain infectious within *T. pyriformis* indicates that virus uptake by protists does not always involve consumption nor trigger immediate energy return to food chains. Not all grazing of viruses by protists thus constitutes virovory. However, uptake by motile protists may lead to the dissemination of infectious viruses in aqueous systems beyond passive transportation. The ecological impact of such a dissemination mechanism remains to be investigated.

## Data availability

All raw data will be made available on Zenodo upon acceptance of the manuscript via https://doi.org/10.5281/zenodo.6901029.

## Declaration of competing interest

There are no conflicts to declare.

## Acknowledgments

This work was supported by the Swiss National Science Foundation (grant number 31003A_182468).

## Footnote

† Electronic supplementary information (ESI):

Table S1: Experimental conditions used in samples analyzed by TEM.

Table S2: Statistical analysis of the variability of HAdV2 removal in replicated experiments using 15 h-starved *T. pyriformis*.

Table S3: qPCR results from experiment with formalin-killed versus live *T. pyriformis* co-incubated with HAdV2.

Table S4: Statistical analysis of digestion experiment.

Figure S1: Experimental design to study egestion and digestion.

Figure S2: Effect of Cytochalasin B on food vacuoles of *T. pyriformis.* Full time course monitoring of food vacuole formation and light microscopy image of carbon-stained food vacuoles from Cyt B treated-*T. pyriformis* versus untreated *T. pyriformis*.

Figure S3: Transmission electron microscopy views of HAdV2 particles in food vacuoles of exposed *T. pyriformis*.

Tomograms: https://doi.org/10.5281/zenodo.6808144

## Notes

https://doi.org/10.5281/zenodo.6808144

https://doi.org/10.5281/zenodo.6901029.

### Competing Interest Statement

The authors have declared no competing interest.

