## Supplementary Material for "Uptake without inactivation of human adenovirus type 2 by *Tetrahymena pyriformis* ciliates"

Running title: Interaction of human adenovirus type 2 with ciliates.

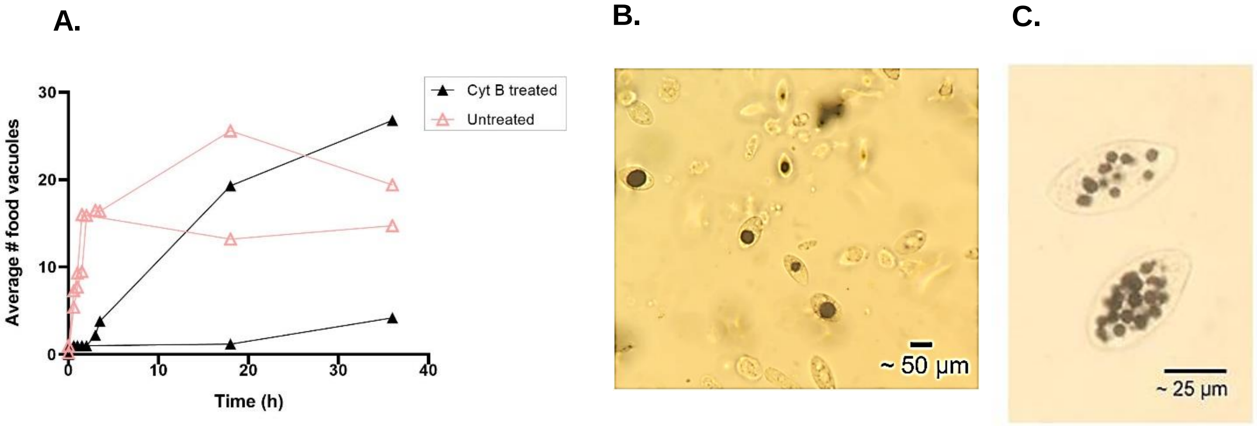


Figure S1: **Effect of Cytochalasin B on food vacuoles (A)** Monitoring of food vacuole formation for Cyt B treated and untreated *T. pyriformis* (full time course) **(B)** Light microscopy image of carbon-stained food vacuoles from Cyt B treated-*T. pyriformis* and **(C)** untreated *T. pyriformis*.

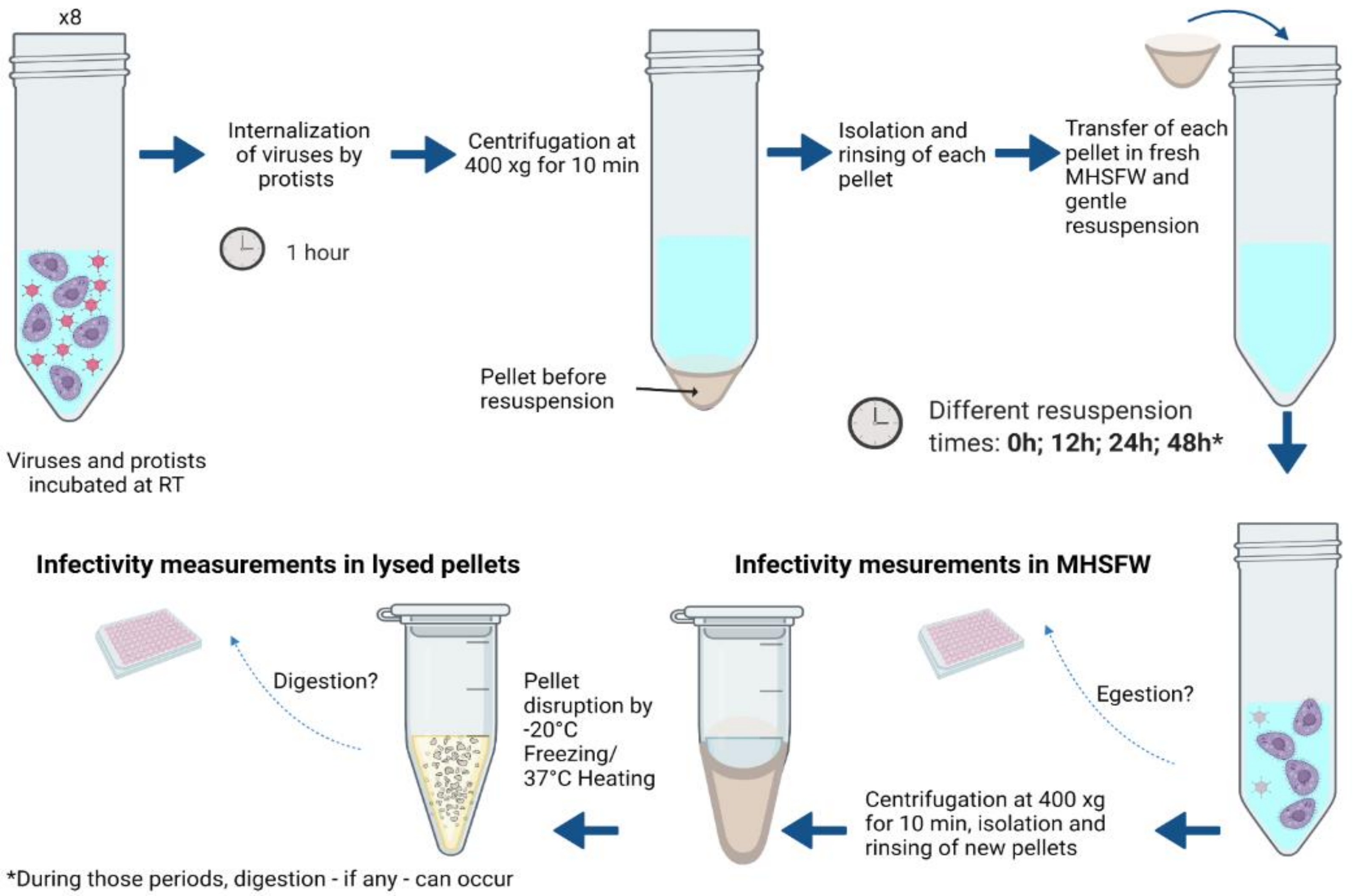


Figure S2: **Experimental design to study egestion and digestion** (created with Biorender).

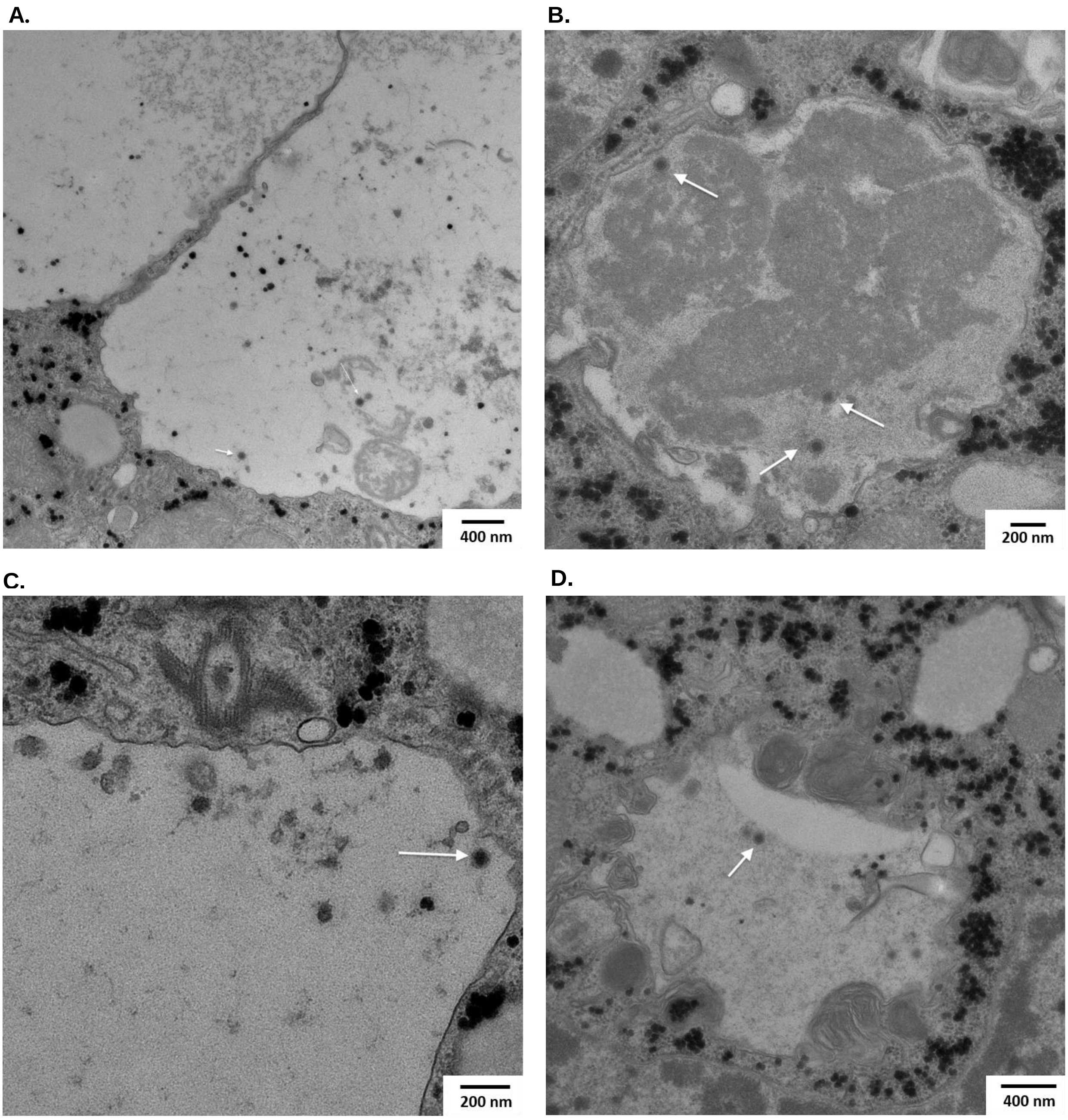


Figure S3: **(A)** to **(D) Transmission electron microscopy views of HAdV2 particles in food vacuoles of exposed *T. pyriformis.***

Table S1: **Statistical analysis of the variability of HAdV2 removal in replicated experiments using 15 h-starved *T. pyriformis*.** After starvation, *T. pyriformis* were co-incubated with HAdV2 for 48 h (Figure 1B). Removal values among individual experiments were compared by ordinary one-way ANOVA with Tukey's multiple comparisons test under the assumption of normal distribution, homogeneity in variances and independent observations.

| **Replicated experiments comparison** | **Mean difference** | **95% confidence interval** | **p-value** |
|---|---|---|---|
| **1 vs. 2** | -0.56 | -1.28 to 0.16 | 0.136 |
| **1 vs. 3** | -0.74 | -1.46 to -0.02 | **0.045** |
| **2 vs. 3** | -0.18 | -0.90 to 0.54 | 0.859 |

Table S2: **qPCR results from experiment with formalin-killed versus live *T. pyriformis* co-incubated with HAdV2.** Corresponding infectivity data is shown in Figure 2. The apparent gain in genome copies in formalin-killed *T. pyriformis* samples after 48 h can be attributed to variability in the qPCR assay.

| | **Live *T. pyriformis*** | **Formalin-killed *T. pyriformis*** | | |
|---|---|---|---|---|
| $\log_{10}(N_{exp} / N_0)$ | -1.8 | 0.7 | 0.8 | 0.8 |

Table S3: **Experimental conditions used in samples analyzed by TEM.**

| Condition | Exposure time to HAdV2 | Initial *T. pyriformis* concentration (cells × $ml^{-1}$) | Initial virus titer (GC × $ml^{-1}$) |
|---|---|---|---|
| A | 50 minutes with respiking | $1 \times 10^5$ | $\sim 1 \times 10^9$ |
| B | 50 minutes with respiking | $3 \times 10^5$ | $\sim 1 \times 10^9$ |

Table S4: **Statistical analysis of digestion experiment** **(Figure 5B).** Virus concentration at different times post transfer to virus-free solution were complared by one-way ANOVA Tukey's multiple comparisons under the assumptions of normal distribution, homogeneity in variances and independent observations.

| Tukey's multiple comparisons test | Mean Difference | 95,00% CI of difference | Adjusted P Value |
|---|---|---|---|
| **0 h vs. 12 h** | -0,050 | -3,312 to 3,212 | 0,982 |
| **0 h vs. 24 h** | -0,60 | -9,298 to 8,098 | 0,630 |
| **0 h vs. 48 h** | -0,80 | -5,149 to 3,549 | 0,281 |
| **12 h vs. 24 h** | -0,55 | -5,986 to 4,886 | 0,475 |
| **12 h vs. 48 h** | -0,75 | -1,837 to 0,337 | 0,077 |
| **24 h vs. 48 h** | -0,20 | -4,549 to 4,149 | 0,787 |
